# An automated interface for sedimentation velocity analysis in SEDFIT

**DOI:** 10.1101/2023.05.14.540690

**Authors:** Peter Schuck, Samuel C. To, Huaying Zhao

## Abstract

Sedimentation velocity analytical ultracentrifugation (SV-AUC) is an indispensable tool for the study of particle size distributions in biopharmaceutical industry, for example, to characterize protein therapeutics and vaccine products. In particular, the diffusion-deconvoluted sedimentation coefficient distribution analysis, in the software SEDFIT, has found widespread applications due to its relatively high resolution and sensitivity. However, a lack of available software compatible with Good Manufacturing Practices (GMP) has hampered the use of SV-AUC in this regulatory environment. To address this, we have created an interface for SEDFIT so that it can serve as an automatically spawned module with controlled data input through command line parameters and output of key results in files. The interface can be integrated in custom GMP compatible software, and in scripts that provide documentation and meta-analyses for replicate or related samples, for example, to streamline analysis of large families of experimental data, such as binding isotherm analyses in the study of protein interactions. To test and demonstrate this approach we provide a MATLAB script mlSEDFIT.

## Introduction

Sedimentation velocity analytical ultracentrifugation (SV-AUC) is a classical first-principles based method to study particles that sediment or float as a result of a gravitational field generated in a centrifuge [1–3]. In the last two decades, this technique has been used in a growing number and types of applications, coinciding with a significant advance in the computational data analysis [4]. Specifically, the combination of efficient Lamm equation modeling [5], novel size-distribution techniques for direct data fitting with and without diffusional deconvolution [6–8], and systematic noise analysis [9,10] as combined in the ls-*g*\*(*s*) and the high-resolution *c*(*s*) approach implemented in the software SEDFIT has garnered widespread applications. These include analyses of size-distributions, hydrodynamic properties, and interactions of particles across the entire size-range accessible to SV-AUC from below 1 kDa to above 10 GDa, ranging from small carbohydrates and peptides [11–13] to proteins and protein complexes [14–17], carbohydrates [18], synthetic polymers [19], small and large nanoparticles [20,21], multi-protein complexes and interacting systems [15,22–24], lipid vesicles and emulsions [25,26], viral particles [27– 32], and entire cellular organisms [33], at concentrations from picomolar to millimolar [3,34,35]. Due to the high resolution and sensitivity, and the measurement of particle sedimentation free in solution in the absence of surfaces or labels, this approach has also proven to be advantageous in biotechnology for the characterization of therapeutic and vaccine products, for example including antibody and other protein therapeutics [36–45], protein/polymer conjugates [46,47], and AAV products [27–31].

One of the challenges of applying SV-AUC in the context of biopharmaceutical production is the lack of software compliant with current good laboratory practices (cGLP) and current good manufacturing practices (cGMP) that satisfies part 11 of Title 21 of the Code of Federal Regulations; Electronic Records; Electronic Signatures (21CFR11) by FDA [48]. Other existing limitations are the required expertise and the laboriousness of serial analyses, for example, of replicate sets or of titration series in the study of interacting systems [41,42,49]. Both would greatly benefit from some degree of automation and streamlining of the data analysis. For this purpose, the present work presents an extension of the SEDFIT software for partially automated operation through the use of a command line interface. The interface allows starting SEDFIT in a specific state, as well as communicating data, analysis parameters, and results. Any secondary software utilizing this interface can spawn SEDFIT, automatically load specific data, execute a specific model for analysis, and retrieve the best-fit results of SEDFIT for documentation, further aggregation, and to carry out meta-analyses of multiple sedimentation coefficient distributions. While this allows streamlined SV-AUC analysis of replicates and titration series, importantly, this approach will be particularly suitable for preserving custody of data and associated analysis results and generating audit trails consistent with 21CFR11.

As illustrated in the present work, the results from such command-line controlled analyses are equivalent to the standard entirely manual use of SEDFIT. Due to the universality of SV-AUC, this approach can be applied to the analysis of therapeutic proteins, carbohydrates, nucleic acids, protein/polymer conjugates, any viral vectors (recombinant or wildtype) including AAV, adenovirus, and lentivirus, lipid vesicles and lipid-based nanoparticles, metal- and other nanoparticles with and without protein conjugates or associated macromolecules such as loaded nucleic acids or proteins.

## Method

The general strategy for command line operated SEDFIT is for it to be spawned by a secondary software, initiating a controlled input of data and providing a controlled return of the analysis results. This will allow logging of all activities and safely record results for further processing. SEDFIT may be also manually executed from the command line, for example, from within the DOS command prompt in Windows, although this would not take advantage of much of the graphical user interface for loading and saving data that is provided in the conventional stand-alone operation of SEDFIT.

The command line controlled run of SEDFIT requires three command line parameters. The first is a number that identifies the SEDFIT mode of operation, at present with possible values of “111” and “112”. The mode “111” will invoke SEDFIT with a standard menu, and “112” will present SEDFIT with a reduced set of menu functions more narrowly tailored to elementary size-distribution analyses. In this restricted mode, non-standard settings are inaccessible to the operator. Different values of this command line parameter are possible in future implementations with alternate behavior. The second command line parameter is an arbitrary string that serves to identify the SEDFIT call. It will be repeated by SEDFIT after completion of the analysis, and thereby provides a layer of security that the reported results do belong to the requested analysis. The third parameter is the path of the input file. For example, the command line could be

~~~
**“sedfit.exe 111 handshakestring c:\datafolder\TestInput.xml”**
~~~

(without the quotes) where ‘handshakestring’ can be replaced by any string, and ‘c:\datafolder\TestInput.xml’ should be replaced by the appropriate directory and filename to point to a valid xml input file in the correct format.

For convenience in generating and parsing the input file, it is in xml format version 1.0, as indicated in the first line by the prolog string `<?xml version=“1.0” encoding=“UTF-8”?>`. This is followed by the root ‘cGMPSedfitCall’ as written in an initial `<cGMPSedfitCall>` call, which is paired with a final `</cGMPSedfitCall>` statement at the end of the file. In between these opening and closing root identifiers are the analysis parameters. They can be in any order, but must adhere to the xml syntax of `<ParameterName>value string</ParameterName>`, where ‘ParameterName’ is any of the identifiers listed below, and ‘value string’ can be a number, a file path, the string `TRUE`, `FALSE`, or other string, as appropriate for the parameter. A complete list of all parameters, their purpose and possible values can be found in **Table 1**. Unless the menu of SEDFIT is restricted (in mode “112”, see above), all parameters can also be set later as usual in the SEDFIT user interface by the operator.

**Table 1.**
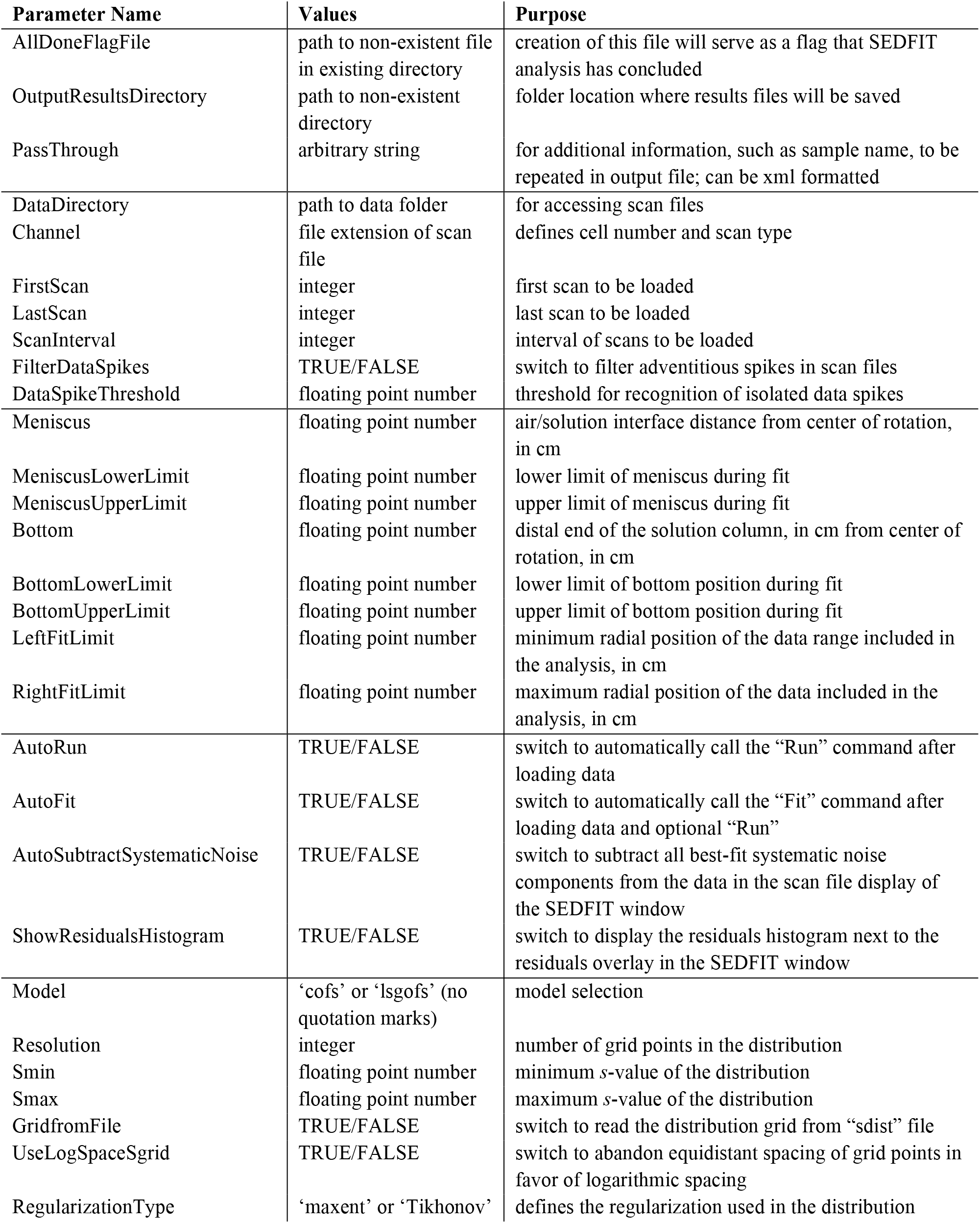
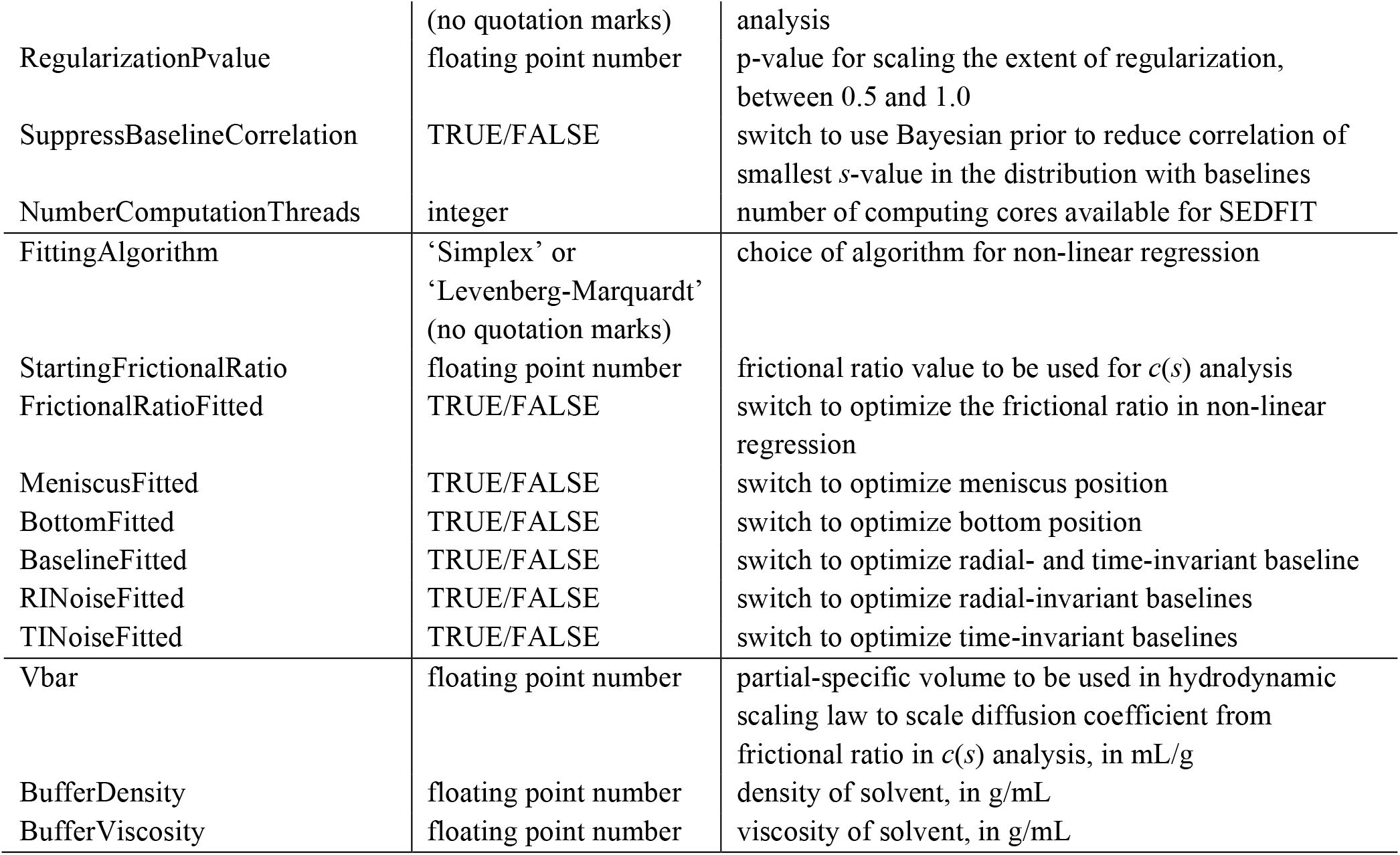
List of Input Parameters. Input parameters will be read from a file named as third command line parameter. It is in xml format and parameters are case sensitive.

While most of the input parameters are strictly related to the data analysis, some are related to the SEDFIT interface and analysis flow. Most importantly, the parameter `AllDoneFlagFile` will establish the path and name of the file that will be created upon completion of the SEDFIT analysis. For example, it might be called ‘c:\datafolder\finished.txt’. Importantly, this file must not exist prior to the SEDFIT call, such that its creation can be used as a convenient flag to the secondary software that SEDFIT has created the results files and was exited. The only content of this file will be the handshake string (second command line parameter, see above), which may be checked by the secondary software for consistency with the calling parameter. The parameter `OutputResultsDirectory` is a path that points to the directory where SEDFIT will create several output files describing the results after conclusion of the analysis, as detailed below. For example, the string ‘c:\datafolder\analysisresults\results1’ will cause SEDFIT to create the subfolder ‘results1’ within the existing ‘c:\datafolder\analysisresults’ directory, and all results files will be located in this ‘results1’ subfolder. Finally, it is possible to use a parameter `PassThrough` to declare any string that, for convenience and clarity, should be repeated in the output xml file. For example, this could be a sample name, user notes, or some additional control strings to be recorded alongside the results. It may also be itself an xml-formatted string, but in this case it should not contain any regular parameter identifiers used in the SEDFIT interface to avoid conflicts.

To define the data to be analyzed, the input file must specify the folder containing the SV-AUC scan files in the parameter `DataDirectory`, the parameter `Channel` that identifies the scan file extension, and the parameters `FirstScan`, `LastScan`, and `ScanInterval` that specify the set of scans to be loaded. It is assumed that the scan files are in the customary SV-AUC numeric format with leading zeros. This information substitutes for manually invoking the Load New Files function in the Data menu of SEDFIT. For example, if `DataDirectory` is ‘c:\datafolder\Run158’, `Channel` is ‘IP2’, `FirstScan` is ‘1’, `LastScan` is ‘101’, and `ScanInterval` is ‘10’, SEDFIT will load 11 files named ‘c:\datafolder\Run158\00001.ip2’, ‘c:\datafolder\Run158\00011.ip2’, ‘c:\datafolder\Run158\00021.ip2’, …, ‘c:\datafolder\Run158\00101.ip2’. The parameters `FilterDataSpikes` and `DataSpikeThreshold` will control whether to ignore isolated spikes in the scan data and the threshold for defining a spike, analogous to the corresponding Loading Options and Tools function of the Options menu of SEDFIT. If the parameters relating to spike filtering are not set then the default filtering with a threshold of 0.4 will be applied.

The input parameters related to the SEDFIT workflow include `AutoRun` and `AutoFit`, both of which take Boolean values `TRUE`or `FALSE`. If true they will automatically initiate the SEDFIT Run or Fit command, respectively, directly after loading the data. This will be equivalent to manually invoking these commands from the SEDFIT menu. (It should be noted that when using `AutoFit` the analysis is carried out before the SEDFIT window appears on the screen.) To enhance the performance of SEDFIT, a parameter `NumberComputationThreads` can be set according to the computational cores that SEDFIT should employ for multithreaded computations. In *c*(*s*) distribution analyses in SEDFIT, key computational steps will be approximately *n*-fold faster if *n* > 1 threads are specified and available [50]. While the default is 2, the optimal value will depend on the computer hardware and on concurrently running software, which may be other instances of SEDFIT. Finally, parameters related to the graphical data presentation are `ShowResidualsHistogram` and `AutoSubtractSystematicNoise`, which will take Boolean values `TRUE`or `FALSE`and control the visual appearance of the SEDFIT analysis window as in their corresponding SEDFIT menu functions [9,10,51]. In addition to changing the graphical display of the SEDFIT window during data analysis, an image of this window will be saved for documentation at the end of the analysis. Scan data, best-fit values, and residuals will also be saved in an output file, so that the graph showing the quality of fit can be recreated (see below).

Ancillary parameters needed for the analysis are the estimated locations of the meniscus and bottom of the solution column, and their fitting limits. `Meniscus` and `Bottom`, as well as `LeftFitLimit` and `RightFitLimit`, are specified as numerical values in cm from the center of rotation, and will function equivalently to their customary graphic input in SEDFIT. During the SEDFIT analysis, they can still be readjusted by the operator or by the fitting routine. If the parameter `MeniscusFitted` is set `TRUE`, the meniscus value will be optimized during the fit, remaining within the bounds specified in the `MeniscusLowerLimit` and `MeniscusUpperLimit` parameters (which should bracket the `Meniscus` value and be smaller than `LeftFitLimit`). Fitting for the meniscus is usually recommended, and therefore coarse initial estimates may suffice, for example, from a table of expected solution column heights dependent on filling volume in standard double-sector cells provided in [3], from a simple identification of the meniscus artifact in scan files, or from a preliminary analysis. Analogous parameters are available for fitting the bottom of the solution column (**Table 1**), however, unless significant back-diffusion is affecting the sedimentation process, typically the bottom position does not need to be fitted in standard analysis and the corresponding value for `BottomFitted` would be `FALSE`.

Most importantly, the parameter `Model` specifies the data analysis model. It currently can take values of ‘lsgofs’ for the ls-g*(s) method [7,8] and ‘cofs’ for the *c*(*s*) method [6,7]. Specifying these models is equivalent to their selection in the Model menu of SEDFIT. Both are sedimentation coefficient distribution models that require definition of the discrete grid of *s*-values through `Smin`, `Smax`, and `Resolution`. Analogous to the manual entry of these parameters in the model parameter boxes of SEDFIT, they take numerical values describing the range of the distribution and the number of grid points. Alternatively, if the parameter `GridfromFile` is set `TRUE`, the grid of s-values can be read from a file ‘sdist’ with the same extension as the scan files (such as ‘sdist.ip2’) that must be located in the same folder as the scan data. This text file will automatically be created after each analysis and contains a single column of *s*-values of the distribution grid, but it can also be edited and augmented and serve as a template for new distribution analyses with custom-spaced grid, for example, to efficiently describe very small or very large sedimenting species outside the range of the majority particles of interest. Another option to modify the grid is `UseLogSpaceSgrid`, which when `TRUE`abandons equidistant grids in favor of logarithmically increasing *s*-value intervals. When using the command line interface these grid functions operate identically with their common stand-alone use of SEDFIT.

The *c*(*s*) analysis applies diffusional deconvolution to achieve sedimentation coefficient distributions with high hydrodynamic resolutions. When using the SEDFIT command line interface, the most commonly used standard *c*(*s*) variant is applied where the diffusion coefficients associated with each sedimentation coefficient are based on a hydrodynamic scaling law *via* a constant hydrodynamic frictional ratio. Thus, the parameter `StartingFrictionalRatio` must be set. The numerical values of the predicted diffusion coefficients also depend on the partial specific volume `Vbar` (to be specified in mL/g) and the buffer density `BufferDensity` (in g/mL). During the analysis, non-linear regression will be used to optimize the frictional ratio during the fit when `FrictionalRatioFitted` is `TRUE`. It should be noted that, unless the final frictional ratio is to be interpreted quantitatively, the partial specific volume values can be rough estimates. The distribution analysis also requires regularization to avoid spurious peaks and error amplification [6,50], which is set through the parameter `RegularizationType`. It currently can take values ‘maxent’ for maximum entropy regularization or ‘Tikhonov’ for Tikhonov-Philips regularization [6]. As is standard in SEDFIT, the tolerated increase in the root-mean-square deviation (rmsd) of the fit allowed for regularization is scaled by F-statistics and a p-value specified in the parameter `RegularizationPvalue` [7]. Since sedimentation patterns of very small particles are very similar to baseline offsets, a correlation between baselines and distribution values at small *s*-values can exist. As usual, this correlation can be suppressed through a Bayesian prior at the smallest *s*-value [52], and this is specified by setting the parameter `SupressBaselineCorrelation` to `TRUE`.

Lastly, details of the fit can be specified through the command line interface. The fitting algorithm is chosen by setting the parameter `FittingAlgorithm` to either ‘Simplex’ or ‘Levenberg-Marquardt’. In addition to the above mentioned fitting of meniscus and bottom of the solution column and the frictional ratio, the treatment of baselines can be specified in the parameters `BaselineFitted`, `RINoiseFitted`, and `TINoiseFitted`. If set to `TRUE`this will cause a spatio-temporally uniform baseline, a time-dependent baseline, and/or a radial-dependent baseline to be fitted during the nonlinear regression, respectively [9,10]. It should be noted that even when the SEDFIT operation is set to automatically execute a Run and Fit command, these procedures can be interrupted and/or manually executed by the SEDFIT operator as usual. Even in the reduced SEDFIT menu it is possible to readjust solution column parameters and other fitting parameters to achieve the best fit prior to concluding the analysis. The usual side effect of the SEDFIT analysis is the creation of the above mentioned ‘sdist’ file containing the *s*-value grid of the distribution analysis, as well as a file named ∼tmppars that, when manually reloading the scan files will allow to restore the last best-fit analysis. Both files are located in the data directory and have the same file extension as the scan files loaded.

When the operator invokes the Exit function of SEDFIT, several files are created prior to the termination of the SEDFIT process and placed in the designated output folder defined in the `OutputResultsDirectory` parameter. The first is a bitmap image of the SEDFIT window, saved as ‘screenshot.bmp’. Besides the graphical display of the scan file overlay and fit, residuals overlay, distribution, and optional residuals bitmap, it shows the customary informational text that provides information about the data files and fitting parameters including the overall rmsd of the fit. Further, it creates text files ‘RInoise.dat’ and ‘TInoise.dat’ which contain two columns with the best-fit time- and radial-dependent baseline values. A file ‘ScanRMSD.dat’ provides information about the rmsd of the fit to each scan file separately. This may help to recognize trends or outliers. The file ‘distribution.dat’ contains two columns with the distribution in form of grid *s*-values *vs. c*(*s*) values. This distribution file can later be integrated used for further analysis in the secondary software. A file ‘dfr.dat’ saves the fitted boundary data for recreation of the data and fit overlay plot, and a residuals plot or bitmap. It is an ASCII text file in the form of a matrix with columns: radius (TI noise), TI noise, RI noise, radius (scan 1), raw data (scan 1), fit value (scan 1), radius (scan 2), raw data (scan 2), fit value (scan 2), etc., and rows corresponding to consecutive radii or scan times, respectively. Finally, SEDFIT creates an output file ‘ResultParameters.xml’ in the same xml format as the input file and containing the same parameters, some of which may have changed due to adjustments during the fit or by operator actions. In addition, it reports the SEDFIT version, file paths of all input SV-AUC scan data files and all output files, as well as the `PassThrough` parameter.

As statistical measures of the quality of fit the output file reports the overall rmsd (`RMSD`), the number of data points fitted (`RMSD-points`), the sum of squared residuals (`RMSD-SSR`), the runs test Z-value (`RunsTestZ`), and the histogram H (`HistogramH`) [7,51]. Additional information about the data include whether scan file time stamps could be accessed for correction (`CheckTimeStamps`) [53], the rotor speed (`RotorSpeed`), the time and accumulated ω^2^t value of the last scan (`tLastScan` and `w2tLastScan`), the rotor temperature at the time of the first and last scan (`TemperatureStart` and `TemperatureEnd`), as well as the temperature average and largest temperature difference (`TemperatureAverage` and `TemperatureDiffMax-Min`). These parameters may be used for experimental quality control to flag the possible presence of convection artifacts [54,55]. A full list of the output parameters can be found in **Table 2**.

**Table 2.** List of Additional Output Parameters. Output will be written in an xml formatted file in the designated output folder. Output parameters include the same parameters regarding data, model, and solution conditions as the input parameters, but also include the additional parameters in this table.

| Parameter Name | Values | Purpose |
| --- | --- | --- |
| SEDFITVersion | string | document the SEDFIT version |
| ScreenShotPath | path to screenshot.bmp file | documented screenshot at end of analysis |
| CofsDataPath | path to distribution.dat file | two-column ASCII text file with the best-fit sedimentation coefficient distribution for further analysis |
| TINoisePath | path to TInoise.dat file | two-column text file with TI noise profile as function of radius to aid in graphical display of scan files |
| RINoisePath | path to RInoise.dat file | two-column text file with RI noise as function of time to aid in graphical display of scan files |
| ScanRMSDpath | path to ScanRMSD.dat file | two-column text file with local rmsd for each scan file, to detect outliers and trends |
| ScanDataFilesLoaded | list of paths | document all the scan files that were fitted |
| PassThrough | arbitrary string | for additional information, as specified in the input parameter file |
| CheckTimeStamps | TRUE/FALSE | for documentation whether time stamps of the scan files were accessible for scan time correction |
| RotorSpeed | integer | rpm of the experiment, as read from scan files |
| Wavelength | floating point number | wavelength entry read from the scan files |
| w2tLastScan | floating point number | $\omega^2t$ entry read from last scan file |
| tLastScan | floating point number | time entry read from last scan file |
| TemperatureStart | floating point number | temperature entry read from first scan, to document temperature discrepancies as a possible source of convective artifacts |
| TemperatureEnd | floating point number | temperature entry read from last scan |
| TemperatureAverage | floating point number | average temperature entries from all scans |
| TemperatureDiffMax-Min | floating point number | range of temperature variation during the SV run |
| FittingStepsText | string | top line of the SEDFIT fitting information display, containing detailed information about the algorithm and steps during non-linear regression |
| RMSD | floating point number | final rmsd of the fit |
| RMSD-points | integer | total number of data points fitted |
| RMSD-SSR | floating point number | sum of squared residuals of the fit |
| RunsTestZ | floating point number | result of the runs test for statistics of runs of positive and negative residuals, as a measure for randomness of residuals |
| HistogramH | floating point number | statistical analysis of the similarity of the residuals histogram to a Gaussian |
| FrictionalRatio | floating point number | best-fit frictional ratio after $c(s)$ analysis |

To indicate termination of the SEDFIT analysis and to allow hand-over to the secondary software, the `AllDoneFlagFile` is created. As mentioned above, this ASCII text file contains as sole entry the handshake string from the command line starting SEDFIT. Creation of this file also indicates that the other results files have been created, which will not be the case if SEDFIT is prematurely exited.

In summary, to access the SEDFIT command line interface any secondary software must carry out the following tasks: 1) organize data such as scan files and starting analysis parameters; 2) generate an input xml file that contains the desired SEDFIT completion flag file (which must not exist yet) and designate the directory for results files; 3) execute SEDFIT with the command line parameters, including a handshake string; 4) wait for completion of the analysis by periodically checking for the creation of the specified completion flag file containing the handshake string; 5) read the results from the xml and other output files created by SEDFIT. 6) Perform optional quality checks, optional integration of the distribution results, carry out secondary analyses, and/or write automated reports. These tasks can be wrapped in access controlled environment, as necessary in the GMP setting. To allow efficient analyses of a large number of samples, the SEDFIT interface can be run in multiple instances side-by-side; if care is taken to create unique `AllDoneFlagFile` files and `OutputResultsDirectory` locations, different SEDFIT instances will operate completely independently of each other.

SEDFIT version 16.50 equipped with the command line interface, as well as the source code of an accompanying family of MATLAB scripts mlSEDFIT can be freely downloaded from sedfitsedphat.nibib.nih.gov/software. Example input and output files can be downloaded from sedfitsedphat.nibib.nih.gov/tools.

## Results

To test the command line interface we wrote a family of MATLAB scripts for data input and retrieval of results, termed ‘mlSEDFIT’. The script may be taken as a template for further modification. We applied it to the analysis of stressed NISTmAb monoclonal antibody [56] that is partially denatured and presents a series of oligomeric populations and a multimodal sedimentation boundary (**Figure 1**). Using the `AutoRun` and `AutoFit` option, the non-linear regression converges at an rmsd of 0.006743 OD (**Figure 1**, solid lines) with a best-fit meniscus at 6.1657 cm, a best-fit frictional ratio of 1.37, and the *c*(*s*) distribution shown in **Figure 2**.

**Figure 1.**
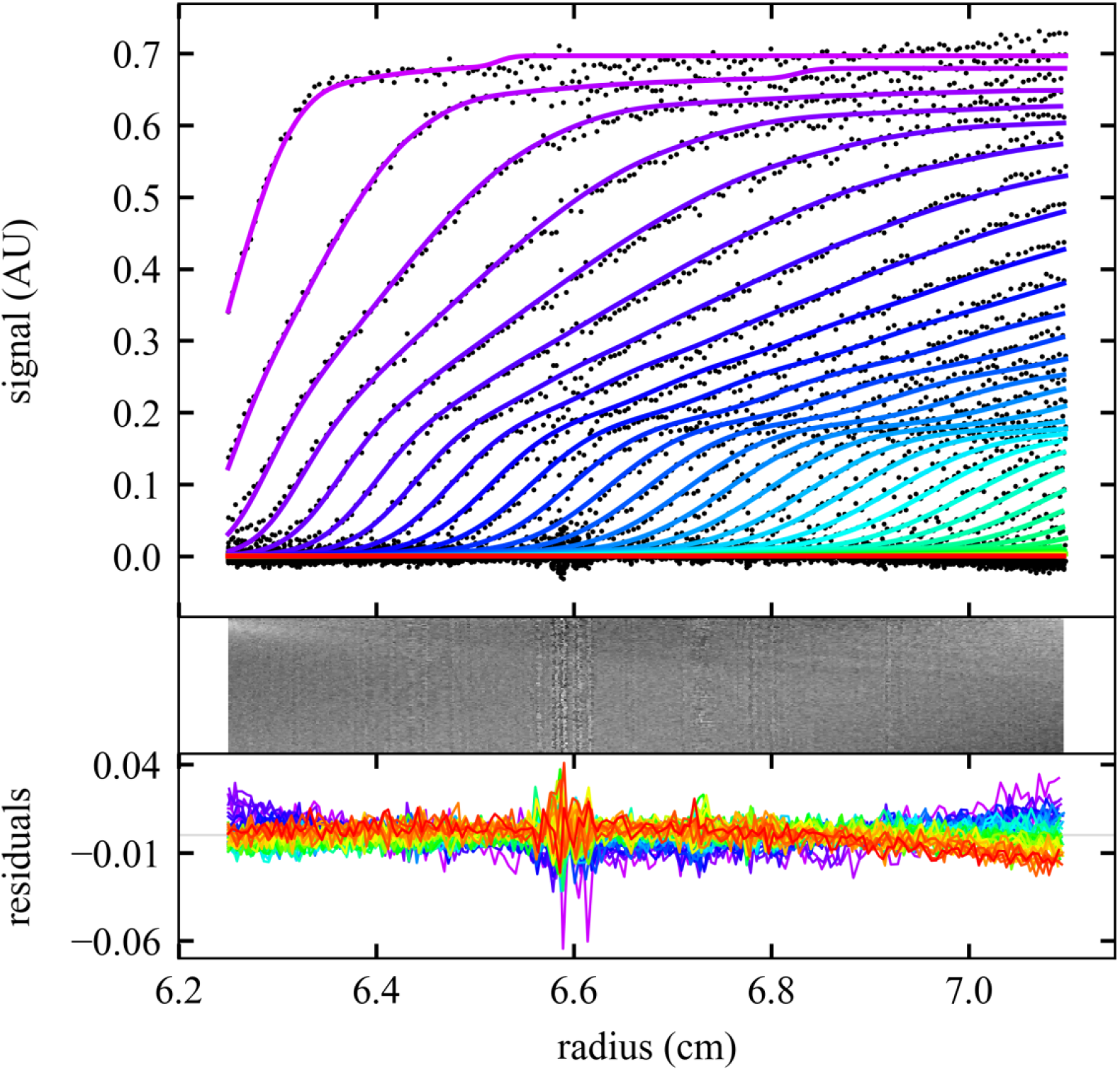
Sedimentation analysis of a NISTmAb sample at 50,000 rpm and 20 °C using the command line operation of SEDFIT. Top: Scan files and best fit (for clarity, showing black dots only for every 2^nd^ data point of every 2^nd^ scan) with a *c*(*s*) model automatically converged to a final rmsd of 0.006743 OD (colored lines). Progression of scan time is indicated by color from purple to red. Middle and Bottom: Residuals bitmap and residuals overlay. Plot was made using the software GUSSI [57], which is spawned from the script mlSEDFIT.

**Figure 2.**
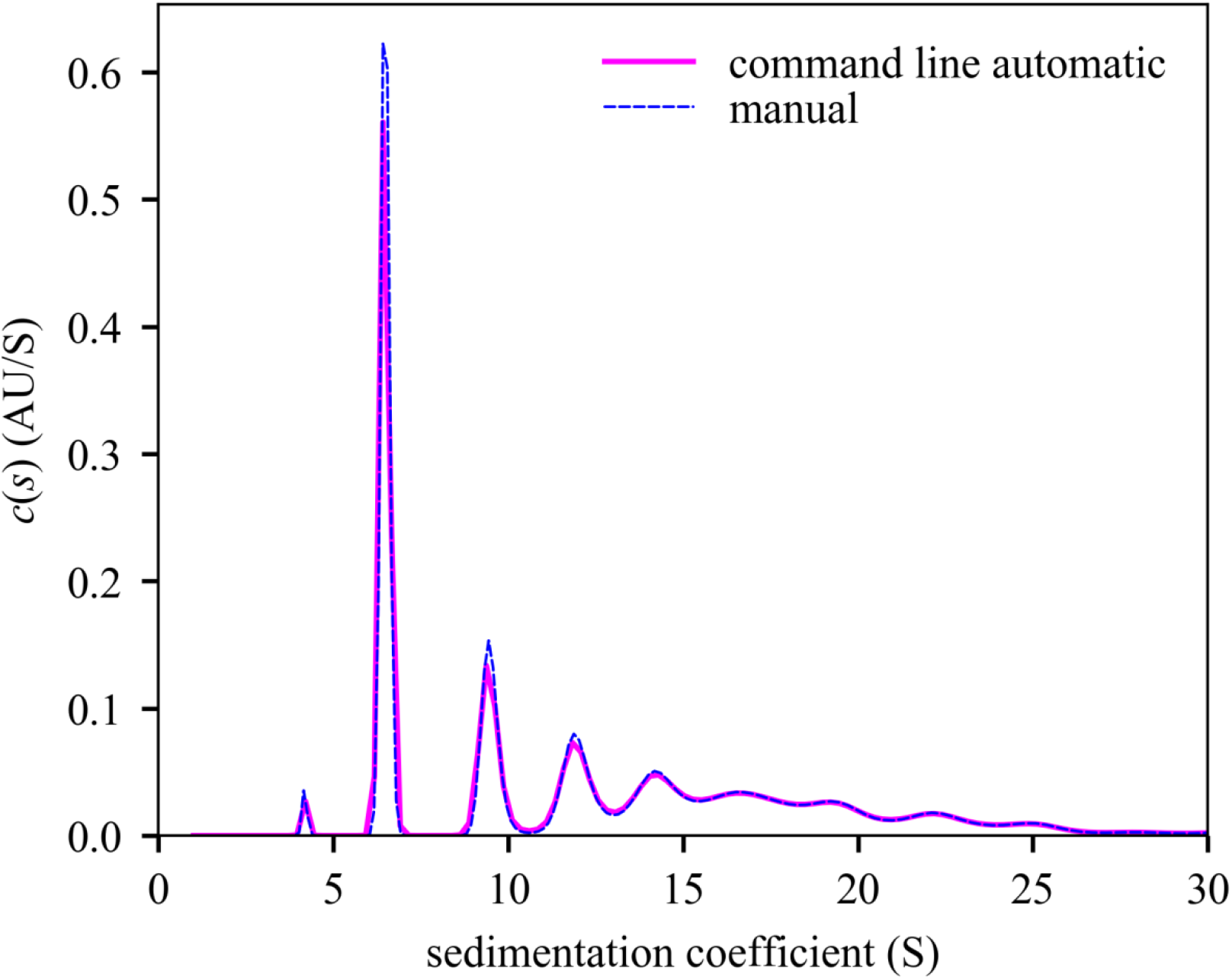
Comparison of *c*(*s*) distributions computed with the command line initialization of SEDFIT and with manual operation. The distribution from command line operation (Figure 1), and exhibits a monomer peak at 6.477 S with 29.198% of signal, a trace degradation product at 4.199 S with 0.95% of signal, a dimer peak at 9.473 S with 12.511% of signal, and higher aggregates with collective s_w_ 16.799 S and 51.774% of signal. The analogous manually operated analysis producing a monomer peak at 6.481 S with 29.274% of signal, a degradation product of 4.178S with 0.922% of the signal, a dimer peak at 9.488 S with 12.511% of signal, and higher aggregates with collective s_w_ of 16.81 S with 51.728% of signal. Integration and plot were made using the software GUSSI [57], which can be spawned from the script mlSEDFIT.

It is possible to manually reload the command line generated analysis and inspect the fit further. However, as an independent control we loaded the same data separately in standard operation of SEDFIT and performed an analysis with the same model. It converged to an rmsd of 0.006738 OD, with a best-fit meniscus at 6.1655 cm, a best-fit frictional ratio of 1.35, and a *c*(*s*) distribution that is virtually identical in all aspects to the distribution from command line operated SEDFIT analysis (**Figure 2**). Differences of < 0.01 S in sedimentation coefficients and < 0.1% in population for all peaks are observed.

Generally, one practical limitation in comparing analysis results can be the required graphical input of the fitting limits, which is replaced by numerical control in the command line parameters. Similarly, detailed loading option preferences may differ in the two operation modes. Furthermore, small numerical differences in repeat analyses should be expected with the Simplex algorithm for fitting, since this involves initial randomization of fitting parameters. Small differences may also be found when adopting different paths in the error surface during non-linear regression. In the present case, remaining insignificant differences in the fit is a result of a locally very flat error surface for the precise numerical value of the frictional ratio parameter, which ultimately reflects the limit of diffusion information content of the broad sedimentation boundaries.

The script mlSEDFIT for testing and demonstrating the new interface is equipped with functions for automatically pre-determining and estimate for the meniscus position for absorbance data prior to spawning SEDFIT, and for creating high-quality illustrations by spawning the software GUSSI [57] utilizing the data, fit, and residuals values retrieved from SEDFIT after the data analysis. In addition, it can integrate the distributions alternatively through a graphical process or through pre-determined integration limits (**Figure 3**). Finally, it will save the results alongside the SEDFIT output parameters. mlSEDFIT can be easily customized and extended, and may be compiled to prevent further modification.

**Figure 3.**
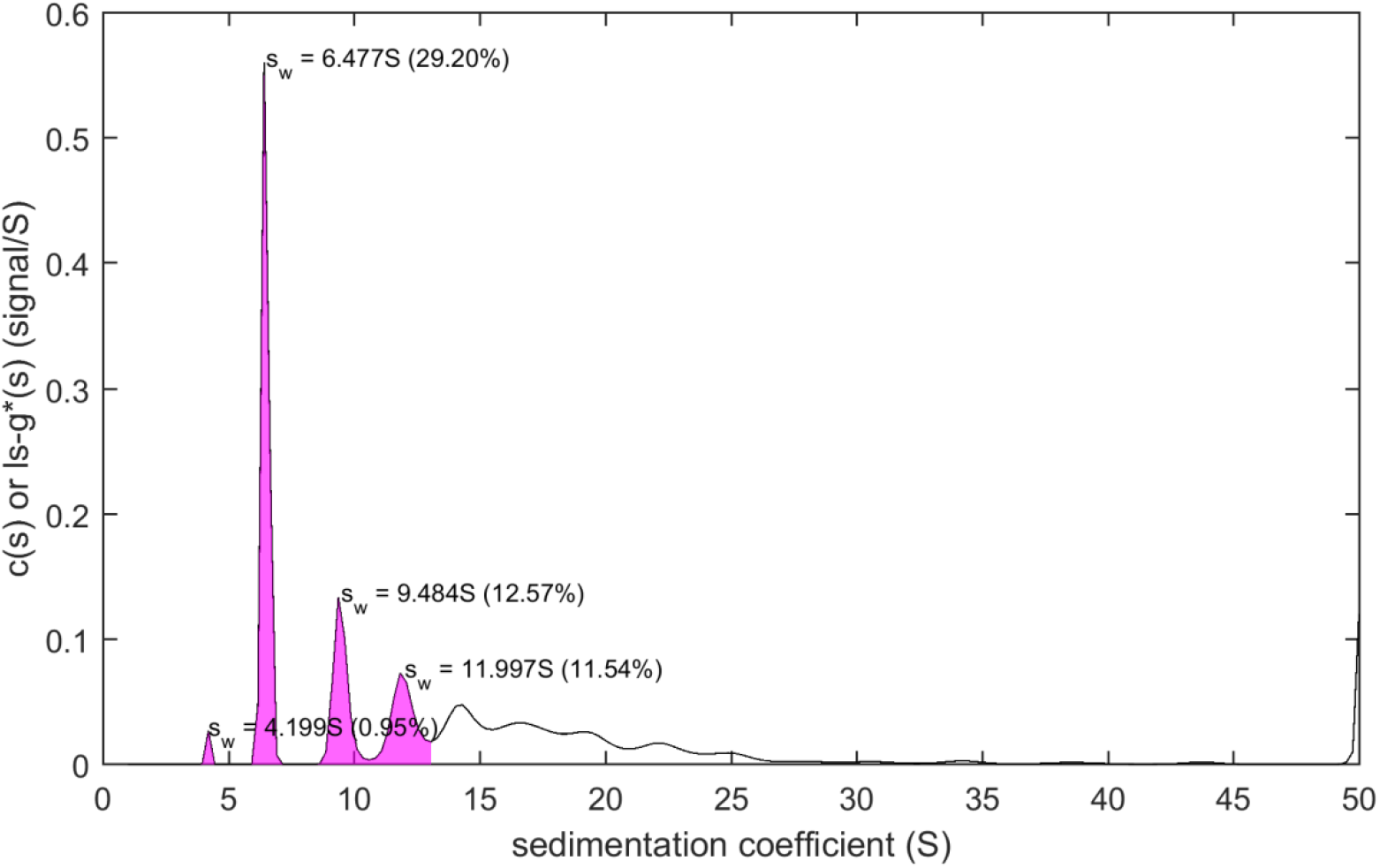
Example for postprocessing of results from SEDFIT analysis in mlSEDFIT. The output generated through the command line interface can be read in the mlSEDFIT script. For example, integration of distribution peaks can be carried out in this script after mouse clicks on the peaks in the distribution plot, as shown.

## Discussion

SV-AUC has become an indispensable tool to study particle size distributions in science and biopharmaceutical industry [36–38,54,58,59]. Therefore, implementation of SV-AUC analyses in the GMP environment would be desirable. The lack of SV-AUC analysis compatible with the GMP environment was previously discussed by Savelyev and colleagues [60]. Their software ULTRASCAN GMP provides data access and analysis workflow control, but unfortunately the SV-AUC data analysis in ULTRASCAN includes *ad hoc* algorithms that are mathematically uncertain in important aspects, and computationally excessively wasteful requiring supercomputers [50]; therefore it has only a minor share in SV-AUC applications in biopharmaceutical applications, where the vast majority of which are carried out with the *c*(*s*) and ls-*g*\*(*s*) methods implemented in SEDFIT [7,39]. Furthermore, ULTRASCAN GMP presents a closed system with pre-conceived workflow strategies and analyses that may not be suitable or adaptable to different objectives. Moreover, it is linked to a particular version of the analytical ultracentrifuge instrument, of which deficiencies for certain applications have been reported [54], and which is currently incapable of providing independent operating system time-stamps of the scan files to the analyst to verify time accuracy [53].

The availability of the computational interface for SEDFIT provides a computational core for flexible state-of-the-art SV-AUC analysis as a module that can be easily embedded into scripts and software satisfying GMP requirements, including auditable analysis trails and custody of data and results, or any other objectives. For demonstration and customization, a generic MATLAB script for spawning SEDFIT was developed, and similar access could conceivably be incorporated in user-friendly software such as GUSSI [57], or in ULTRASCAN GMP [60] or other custom-written GMP software.

At present, the analysis still needs to be supervised, since manual adjustments to the fitting parameters and model may be required for arriving at the best-fit analysis. This allows adventitious scan files or other possible artifacts from experimental imperfections to be recognized and their effect to be alleviated.

Detailed protocols and instructions can be found in the literature [3,7,39,45,61–63], and for reliable results this guidance is equally valid when using the command line interface. Nonetheless, the interface described here can provide a platform for future improvements that conceivably may allow fully unsupervised analyses, for example, of replicate experiments. For series of equivalent experiments, the present version already allows the result (for example, meniscus position or frictional ratio) of an initial analysis to be automatically entered as starting parameters for a following data set, thereby improving the efficiency of the analysis. Besides the GMP environment, this may be useful for analyzing large families of experimental data sets designated for meta-analyses, such as collective integration of distributions for binding isotherms and their analysis [23,49,63]. This ties in with developments of higher throughput experimental techniques, such as pseudo-absorbance data acquisition without need of a reference sector [64] and 3D-printed multi-sector centerpieces [65,66].

Importantly, since none of the computational functions from SEDFIT have been altered, the results will remain the same as in the equivalent standard operation of SEDFIT. The command line interface solely modifies the data input and output, replacing manual startup and loading of analysis files with automated pre-loaded SEDFIT. For this reason, the command line mode of SEDFIT will be applicable to the same range of current and future applications. With regard to the biopharmaceutical industry this includes studies of therapeutic peptides and proteins, polymer conjugates, nucleic acids, carbohydrates, vectors for therapeutics or vaccines based on metal nanoparticles, lipid nanoparticles, viral vectors such as adenovirus, AAV or lentivirus, and others. More generally, due to the universal nature of buoyant mass-based separation in SV-AUC and the high sensitivity and hydrodynamic resolution of *c*(*s*) analysis in SEDFIT, it will be applicable to study mass- and size-distributions of macromolecules and particles that differ in density from that of the formulation buffer across a mass range from 1 kDa to >10 GDa, or a sedimentation coefficient range between 0.1 and 100,000 S [67,68].

## Acknowledgement

This work was supported by the Intramural Research Program of the National Institute of Biomedical Imaging and Bioengineering, National Institutes of Health.

